# Histone H3K9 methylation and Heterochromatin Protein 1 do not limit DNA accessibility in living *S. pombe* cells

**DOI:** 10.64898/2025.12.12.694020

**Authors:** Lalita Panigrahi, Zhuwei Xu, David J. Clark

## Abstract

Chromatin is intrinsically repressive, limiting access to DNA, implying a major regulatory role. Studies with nuclei support this model. However, we have shown previously that genomic DNA is completely accessible in living budding yeast and human cells, except for centromeric chromatin. The fission yeast, *Schizosaccharomyces pombe*, provides a tractable model for heterochromatin. As in mammalian cells, *S. pombe* heterochromatin is marked by histone H3K9 di- and tri-methylation (H3K9me2/3), introduced by the Clr4 histone methylase, and heterochromatin protein 1 (HP1/Swi6). Here, we developed a copper-inducible DNA methyltransferase system to measure heterochromatin accessibility in living *S. pombe* cells. We find that euchromatin and heterochromatin are similarly and generally accessible in vivo, indicating that *S. pombe* chromatin is globally dynamic. *S. pombe* centromeres are also fully accessible, unlike budding yeast and human centromeres. In contrast to living cells, *S. pombe* chromatin is mostly inaccessible in isolated nuclei, primarily due to tight nucleosome spacing with very little linker DNA. Loss of Clr4 or Swi6 has little effect on genome accessibility in live cells or nuclei, suggesting that the *S. pombe* H3K9me/HP1 system does not repress transcription by preventing access to DNA.

## Introduction

Chromatin is the nucleoprotein complex that organizes and packages eukaryotic genomes. Its fundamental repeating unit is the nucleosome, composed of ∼147 bp of DNA wrapped around an octamer of histone proteins composed of two copies each of H2A, H2B, H3 and H4 (1). Nucleosomes are regularly spaced along the DNA, connected by linker DNA, creating the characteristic “beads on a string” appearance when unfolded in the electron microscope. These nucleosomal arrays fold into higher order structures under physiological conditions (2,3). Current models propose that chromatin is intrinsically repressive, potentially restricting access to the DNA at both the nucleosome level and in higher order structures, including heterochromatin (4,5). The degree to which genomic DNA is accessible inside cells is therefore of central importance to understanding gene regulation.

Chromatin structure and function are regulated through histone modifications, incorporation of histone variants, DNA methylation and ATP-dependent chromatin remodeling. Specific histone modifications mark euchromatin and heterochromatin states. For example, acetylation marks such as H3K9ac and H3K14ac are enriched at active promoters and enhancers, whereas methylation marks such as H3K4me3 and H3K36me3 are associated with actively transcribed genes (reviewed in (6)). In contrast, repressed genes in facultative heterochromatin are marked by H3K27me3, whereas constitutive heterochromatin is marked by H3K9me2/3 (reviewed in (7)).

Heterochromatin protein 1 (HP1) is a major component of heterochromatin and occurs in three mammalian isoforms (HP1α, HP1β and HP1γ) (reviewed in (8)). HP1 binds reversibly to H3K9me3, resulting in a dynamic chromatin structure (9–11). Cryo-EM studies have shown that a human HP1 dimer can bridge two H3K9me3-modified nucleosomes without engaging the linker DNA (12). HP1α can oligomerise and form condensates, promoting long-range interactions in chromatin, resulting in compaction (13,14). HP1α also binds H2B independently of H3K9me and promotes unwrapping of nucleosomal terminal DNA (15). These observations suggest a role for HP1 in condensing mammalian heterochromatin. The dynamic nature of HP1 binding suggests that access to the underlying nucleosomal array is not necessarily restricted in heterochromatin. Consistent with this finding, our recent studies in human MCF7 and MCF10A cells, using a sequence-specific DNA methylase as a probe, revealed that nucleosomal DNA is globally accessible in living cells (16). Both facultative heterochromatin and constitutive heterochromatin are accessible in vivo, although they are methylated more slowly than euchromatin, perhaps reflecting compaction (16).

Constitutive heterochromatin is primarily formed on highly repetitive elements (reviewed in (17)) and is therefore difficult to study in mammals. Unlike budding yeast, the fission yeast (*Schizosaccharomyces pombe*) possesses both H3K9me3 and HP1 orthologues (Swi6 and Chp2). *S. pombe* heterochromatin is very well-studied and provides a tractable model system that is much simpler than mammalian heterochromatin. *S. pombe* heterochromatin is located at pericentromeres, sub-telomeres and the mating type locus (18–20). Foreign genes inserted into these regions undergo transcriptional silencing (21,22). The formation of *S. pombe* heterochromatin requires transcription factors (23), RNA polymerase II transcription, the RNAi machinery and histone-modifying enzymes (20). H3K9 methylation is catalysed by Clr4, the sole H3K9 methyltransferase in *S. pombe* (reviewed in (24)). The ability of HP1 to oligomerise is essential for the maintenance of *S. pombe* heterochromatin (Seman 2023). Loss of heterochromatin in a *clr4* null mutant results in altered long range chromosome interactions, suggesting that heterochromatin influences euchromatin architecture (25).

As in higher eukaryotes, *S. pombe* HP1 (Swi6) binds dynamically to H3K9me2/3 (20,26–28). However, *S. pombe* heterochromatin differs from mammalian heterochromatin in that it is not condensed in interphase cells (29). The most condensed chromatin in *S. pombe* cells (termed “knob” chromatin) is located adjacent to sub-telomeric heterochromatin, but lacks HP1 and H3K9me2/3 (29). These data imply that HP1 and H3K9me2/3 are insufficient to compact heterochromatin in vivo. Thus, mammalian heterochromatin may contain additional factors that drive condensation, such as linker histone (H1), which is absent in *S. pombe*.

Nevertheless, *S. pombe* shares many chromatin features with higher eukaryotes. Genome-wide nucleosome mapping has revealed conserved promoter features, including a nucleosome-depleted region (NDR) upstream of the transcription start site (TSS) at most genes, and phased nucleosomes downstream of the TSS at all but the most inactive genes (30,31). The average nucleosome spacing in *S. pombe* is unusually short (∼152 bp), such that there is very little linker DNA, consistent with the absence of H1 (30–32). *S. pombe* centromeres (*cen1, cen2* and *cen3*) span ∼35 kb, ∼55 kb and ∼110 kb, respectively. Each centromere consists of a central core (*cnt*) flanked by innermost repeats (*imr*), which are in turn flanked by heterochromatic *dg* and *dh* repetitive elements (reviewed in (22)).

Here, we have measured the accessibility of the genome in living *S. pombe* cells. We developed a copper-inducible system to express the bacterial DNA adenine methyltransferase (’Dam’) in *S. pombe*. Dam methylates adenines in GATC motifs and serves as a proxy for sequence-specific transcription factors. Using our quantitative DNA accessibility sequencing assay (qDA-seq) (33,34), we find that the *S. pombe* genome is globally accessible in vivo. Strikingly, unlike human heterochromatin, *S. pombe* heterochromatin is as accessible as euchromatin, indicating that both euchromatin and heterochromatin have highly dynamic chromatin structures that are unlikely to present a major barrier to DNA access. Furthermore, loss of Clr4 or Swi6 results in only minor changes in DNA accessibility in vivo.

## Materials and methods

### Plasmids

Plasmid p856 (7652 bp) for integrating the Dam cassette at the *urg1* locus was constructed as follows: A synthetic NotI fragment with the *S. pombe* uracil-inducible *urg1* promoter driving the budding yeast *ACE1* open reading frame (ORF) without a stop codon, followed by PacI and SacI sites and 300 bp of 3’-*urg1*, was inserted into pMK-RQ to obtain p850 (Thermo-Fisher GeneArt). p856 was constructed by insertion of the 3685-bp PacI-SacI myc7-3’*ADH1*-*tCUP1promoter*-Dam-degron-3HA-KanMX fragment from p840 (34). Plasmid p964 (8068 bp) for integrating the M.SssI cassette at the *urg1* locus was constructed by replacing the Dam ORF in p856 with the M.SssI ORF as follows: the 2605-bp BsrGI-ScaI Dam fragment in p856 was replaced with the 2929-bp BsrGI-ScaI M.SssI fragment from p906 (34) to obtain p960. The 215-bp AvrII-SwaI truncated *CUP1* (*tCUP1*) promoter in p960 was replaced with the 310-bp AvrII-SwaI fragment containing the full-length *CUP1* promoter from p834 (34) to obtain p964. Plasmid p951 (9898 bp) for integrating the OsTIR1 gene carrying the F74A mutation was constructed by insertion of a 1701-bp NheI fragment containing a hygromycin resistance (Hph) gene obtained by PCR, using primers 2422 (5’-GACGTTGCTAGCCAGATCTGTTTAGCTTGCCTC) and 2424 (5’-GACGTTGCTAGCGAGCTCGTTAAAGCCTTCGA) with p846 as template, into the NheI site in Padh1-OsTIR1(F74A)-NLS-TADH1-Sp.arg3+/pUC19* (Addgene 169353), oriented such that the Hph and TIR1 genes are transcribed in the same direction. p846 was obtained by replacing the 1451-bp AscI-SacI KanMX gene in p840 with the 1677-bp AscI-SacI Hph gene fragment from pYM24 (Euroscarf P30236). We note that plasmids containing the M.SssI gene isolated from *E. coli* were partially resistant to digestion by restriction enzymes sensitive to CG methylation. Plasmid p998 for tagging Gcn5 at the C-terminus was constructed as follows: A synthetic NotI fragment with the 3’-end of the *gcn5* ORF fused to 3 HA tags (identical in sequence to those fused to Dam in p856), a polylinker including SacI and AscI sites and downstream *gcn5* flanking sequence, was inserted into pMA to obtain p990 (Thermo-Fisher GeneArt). A 1290-bp SacI-AscI fragment containing the NatNT2 gene derived from pYM21 (Euroscarf P30307) was inserted into p990 to obtain p998. The same NatNT2 gene fragment was inserted into synthetic plasmids p987 and p988 (Thermo-Fisher GeneArt) to obtain integration plasmids for deleting the *clr4* open reading frame (ORF) (p995) or the *swi6* ORF (p996) using a NotI digest, such that NatNT2 is in the opposite orientation to *clr4* or *swi6*. p987 contains a NotI insert with 241 bp of 5’-*clr4* DNA and 243 bp of 3’-*clr4* DNA separated by a polylinker including a SacI site and an AscI site. p988 contains a NotI insert with 240 bp of 5’-*swi6* DNA and 236 bp of 3’-*swi6* DNA separated by a polylinker including a SacI site and an AscI site. All plasmids and their verified sequences are available upon request.

### Strain construction

The *S. pombe* strains used in this study are listed in Supplementary Table S1. Strains were prepared using standard genetic methods (35). SP1L was obtained by transformation of wild type strain 972h-(ATCC 24843) with the Dam gene-containing NotI fragment from p856 and selection with G-418. SP3L was obtained by transformation of SP1L with the TIR1(F74A)-containing AleI.v2-HindIII fragment from p951 and selection with hygromycin. SP12L was obtained by transformation of SPT999a (36) with the same TIR1(F74A) fragment from p951 and selection with hygromycin. SP15L was obtained by transformation of SP12L with the M.SssI gene-containing BmrI-HpaI fragment from p964 and selection with G-418. SP27L was obtained by transformation of SP3L with the NotI fragment from p998 containing the *gcn5*-3HA construct, and selection with nourseothricin. Integration at targeted loci was confirmed by PCR. SP23L and SP24L were obtained by transformation of SP3L with the NotI fragments from p995 and p996 containing the *clr4::NatNT2*, and *swi6*::NatNT2 constructs, respectively, followed by selection with nourseothricin. For nuclei experiments, strain 972h-was transformed with the same NotI fragments to obtain SP32L and SP33L, respectively. The deletions were confirmed by PCR.

### Induction time courses for Dam and M.SssI

SP3L, SP15L, SP23L and SP24L were cultured in complete medium (0.5% yeast extract, 3% D-glucose, 0.2 g/l adenine) supplemented with 100 nM 5a-IAA (TCI A3390) at 25°C to mid-log phase (OD_600_ of 0.6 to 0.7). At time zero, samples were collected for genomic DNA extraction (∼35 OD units) and immunoblotting (∼10 OD units) and stored at -80°C. The remaining cells were washed twice with fresh medium by filtration to remove residual 5a-IAA and resuspended in the original volume of fresh medium (0.5% yeast extract, 3% D-glucose, 0.2 g/l adenine, 0.2 g/l uracil). Copper sulphate was added to a final concentration of 0.4 mM for Dam expression (SP3L, SP23L and SP24L) or 1 mM for M.SssI expression (SP15L), and the culture was warmed to 30°C. Cells were harvested 30, 60, 120 and 240 min after copper addition for immunoblotting and genomic DNA as above. For G1 arrest, cells were cultured in complete medium to ∼0.3 to 0.4 x10^7^ cells/ml. The cells were washed twice with EMM No nitrogen (MP Biochemicals 4110712) medium, resuspended in an equal volume of EMM No nitrogen medium with 100 nM 5a-IAA and incubated at 25°C for 8 h to induce G1 arrest by nitrogen starvation. Arrest was verified microscopically, indicated by the characteristic shift from elongated morphology to rounded cell shape. After confirmation of G1 arrest, the cells were washed thoroughly to eliminate 5a-IAA. To initiate Dam expression, the cells were warmed to 30°C and CuSO_4_ was added to 1 mM. Samples were collected at 30, 60, 120 and 240 min after Cu addition.

### Genomic DNA extraction

Genomic DNA was purified by washing cell pellets 3 times with 50 mM Tris-HCl pH 8.0, 5 mM Na-EDTA (5x TE) buffer with 2% SDS to extract cell contents while retaining the cell wall, followed by additional washes with 5x TE to remove residual detergent. The pellets were resuspended in 500 µl 5x TE containing 30 mM 2-mercaptoethanol and 50 µl lyticase (Sigma L2524 at 25,000 U/ml), and incubated at 37°C for 30 min. Cell wall digestion was monitored by measuring OD_600_ in 1% SDS. Subsequently 50 µl 20% SDS was added and the mixture was incubated at 65°C for 5 min. Then 150 µl 5 M potassium acetate was added, followed by an equal volume of chloroform:isoamyl alcohol (24:1). The mixture was shaken vigorously, centrifuged and subjected to a second chloroform extraction. Genomic DNA in the aqueous supernatant was precipitated with 0.7 vol. isopropanol and the pellet was washed with 70% ethanol. The purified DNA was dissolved in 10 mM Tris-HCl pH 8.0, 0.1 mM Na-EDTA and treated with 0.4 mg/ml RNase A (Thermo-Fisher EN0531) for 1.5 h at 37°C. Finally genomic DNA was purified using 0.8 vol. AMPure XP beads (Beckman-Coulter A63881).

### Illumina paired-end library preparation

Genomic DNA was digested with 10 U DpnI (New England Biolabs (NEB) R0176L) at 37°C for 1.5 h. The reaction was terminated by adding 10 mM EDTA, 0.1% SDS. DpnI-digested DNA was purified using 1.8 vol. AMPure XP beads and eluted in 50 µl 10 mM Tris-HCl pH 8.0, 0.1 mM Na-EDTA. DNA was analyzed in a 1% (w/v) agarose gel stained with SYBR Gold (Invitrogen S11494) with MassRuler as a marker (Thermo Scientific SM0403). Any nicks were sealed by treatment with HiFi Taq DNA ligase (NEB M0647S) at 37°C for 1 h, the reaction was quenched with 20 mM EDTA, and the DNA was purified using 1.8 vol. AMPure XP beads as above. The DNA was fragmented using a Covaris ME 220 ultrasonicator in a Covaris tube (Covaris 520166) at 4°C using the 350-bp program with the following settings: peak power 50 W, duty factor 10%, 1000 cycles per burst, average power 5 with a total processing time of 150 s per tube. The DNA fragment size was assessed by electrophoresis in a 2% (w/v) agarose gel stained with SYBR GOLD (Invitrogen S11494). The sonicated DNA was purified using 1.8 vol. AMPure XP beads and eluted in 50 µl 10 mM Tris-HCl pH 8.0, 0.1 mM Na-EDTA. Paired-end libraries were prepared as described (34). Sequencing was performed using an Illumina NextSeq-2000.

### Nuclei isolation and Dam methylation in vitro

*S. pombe* 972h-, SP32L and SP33L cells were cultured in complete medium to mid-log phase, harvested and stored at -80℃. For spheroplast preparation, ∼100 OD_600_ units were resuspended in 15 ml SM buffer (5% yeast nitrogen base, 2% glucose, 1 M sorbitol, 50 mM Tris-HCl pH 8.0, 20 mM 2-mercaptoethanol). Cells were incubated with 25,000 units lyticase (Sigma L2524) at 30°C for 30 min. Digestion was monitored by measuring the OD_600_ of 30 µl cells suspended in 1 ml 1% SDS, and was considered complete when the OD_600_ was < 10% of the initial value. Spheroplasts were collected by centrifugation (8140 g, 5 min, 4°C) and washed once with ice-cold 1 M sorbitol, 50 mM Tris-HCl pH 8.0. Spheroplasts were lysed in 20 ml F buffer (18% Ficoll, 40 mM potassium phosphate pH 6.5, 1 mM MgCl_2_) supplemented with protease inhibitors (Roche 05056489001), 5 mM 2-mercaptoethanol, 2 mM EDTA and 1% Triton X-100, and incubated for 5 min at room temperature. The lysate was layered onto a 15 ml FG buffer step (7% Ficoll, 20% glycerol, 40 mM potassium phosphate pH 6.5, 1 mM MgCl_2_, 2 mM EDTA, protease inhibitors, 5 mM 2-mercaptoethanol) and centrifuged (22,620 g, 20 min, 4°C). The nuclei pellet was resuspended in 2 ml 10 mM HEPES pH 7.4, 100 mM KCl, 5 mM MgCl_2_, 0.2 mM S-adenosylmethionine. Methylation reactions were performed by adding 0, 25, 50, 100 or 200 U Dam methyltransferase (NEB M0222L at 8 U/µl; 1.6 µg/ml) to 100 µl aliquots of nuclei (final Dam concentrations: 0, 1.5, 2.9, 5.6 and 10.0 nM) and incubating at 25°C for 30 min. Reactions were terminated with 0.1 M Na-EDTA, 10% SDS to final concentrations of 10 mM EDTA and 1% SDS. Genomic DNA was purified, digested with DpnI and subjected to Illumina paired-end sequencing, as described above.

### Nanopore sequencing

Genomic DNA was extracted as described above and purified using the PureLink Genomic DNA Mini Kit (Invitrogen K182002). DNA ends were repaired for adaptor ligation using the NEBNext Companion Module for Oxford Nanopore Technologies (ONT) Ligation Sequencing (NEB E7180L). Barcoding and adaptor ligation were then performed using the ONT Native Barcoding kit (SQK-NBD114.24). Sequencing was performed using an ONT MinION flow cell according to the manufacturer’s instructions. Reads were base-called using Dorado v1.0.0 with the dna_r10.4.1_e8.2_400bps_sup v5.2.0 basecalling model and 5mCG_5hmCG v1 modification model (--min-qscore 8). Reads were aligned to both the Pombase reference genome and our de novo assembly for SPT999a (see below) using the Dorado aligner. We filtered reads with at least one 5mCG site (mCG threshold > 0.9 and unmethylated CG < 0.1). The 5mCpG sites from filtered bam files were called using modkit v0.5.0 (-- cpg --combine-strands --ignore h).

### Immunoblotting

Cell pellets (∼10 OD_600_ units) were resuspended in 100 µl LDS sample buffer (Invitrogen NP0007) containing 0.2 M 2-mercaptoethanol and heated at 99°C for 20 min. The samples were briefly centrifuged and the supernatants were transferred to fresh tubes. Proteins were separated in 4-12% Bis-Tris gels (Invitrogen NP0336) using MOPS SDS running buffer (Invitrogen NP001). One gel was stained with Coomassie blue, while the proteins in the other gel were transferred to a PVDF membrane (Invitrogen IB401002) using an iBlot transfer machine according to the manufacturer’s instructions. The membrane was blocked with 5% skimmed milk in TBS (20 mM Tris-HCl, pH 8.0, 0.5 M NaCl) containing 0.1% Tween-20 at room temperature. For HA detection, the membrane was incubated overnight at 4°C on a tube rotator with HRP-conjugated anti-HA antibody (3F10; Roche 12013819001), diluted 1:2500. The membrane was washed 3 times for 10 min each with TBS containing 0.1% Tween-20 at room temperature. Chemiluminescence was detected using the SuperSignal West Pico Plus Chemiluminescent kit (Thermo-Fisher 34577) and visualized using an Azure 600 Chemiluminescent Western blot Imager. The loading control was detected by treating the membrane with stripping buffer (Thermo-Fisher 46430) for 3 min at room temperature on a tube rotator, then washed 3 times with PBST (137 mM NaCl, 2.7 mM KCl, 8 mM Na_2_HPO_4_, 2 mM KH_2_PO_4_, pH 7.4, 0.1% Tween-20). The membrane was re-blocked with 5% skimmed milk in PBST for 1 h at room temperature followed by incubation with HRP-conjugated anti-tubulin antibody (Abcam ab-185067) diluted 1:10000 in PBST for 1 h. After 3 washes with PBST, the signal was developed as described above.

### Computational analysis

Methylated fractions for GATC sites were calculated using the SnakemakeMethylFrac workflow (16). Our decaying sine wave model was used to analyze average phasing data (37). We performed de novo assembly of the chromosome sequences for the SPT999a strain with nanopore sequencing reads using canu/2.2 (38) with parameters: genomeSize=13m, maxInputCoverage=300, rawErrorRate=0.03, correctedErrorRate=0.01, corOutCoverage=100, minReadLength=10000. We included the mitochondria sequence from the Pombase reference genome. The rDNA locus in the reference genome was annotated based on the Pol I initiation site (39) and the BoxI terminator (40). The *dg* and *dh* elements are annotated by the *dg* and *dh* elements in the left pericentromeric region of chromosome I using MUMMER/4.0.1 (41). The mating locus was annotated using liftoff v1.6.3 (42). For H3K9me2/3 peaks, we used ChIP-seq data from GSE280066 (43), which were pre-processed by fastp/0.24.0 (44), and aligned to the SPT999a genome using Bowtie2 v2.5.3 (-X 5000 --very-sensitive --no-discordant --no-mixed --no-unal) (45). Aligned reads were filtered using SAMtools v1.21 (-f 0x2 -F 0x300 -q 1) (46). H3K9me2 and H3K9me3 peaks were predicted using macs v3.0.0 (47); the peaks from their three replicates were combined to annotate H3K9me2/3 regions. Protein expression levels were obtained from Pombase (48) using data for vegetative growth (49,50). RNAseq data were obtained from (51) (GSE227184) and analysed using salmon v2.5.0 (52). The Mann-Whitney U test was used for boxplot comparisons. For boxplots with multiple replicates, data were pooled by group prior to statistical testing. Groups were considered significantly changed if p < 0.05 and effect size (absolute rank-biserial correlation, r) > 0.2.

## Results

### A copper-inducible protein expression system for *S. pombe*

We used the Dam DNA methyltransferase as a probe to measure genome accessibility in living *S. pombe* cells. Dam methylates the adenine in the sequence GATC, if it is accessible. We developed a robust copper-inducible expression system based on the *S. cerevisiae CUP1* promoter. The Dam gene was placed under the control of a truncated *CUP1* promoter in which only one of the two upstream activating sequences (UASs) is present in order to reduce the Dam expression level. In the presence of copper, the DNA-binding domain of the *S. cerevisiae* Ace1 transcription factor folds and binds specifically to the *CUP1* UAS to activate transcription (53,54). The *ACE1* gene was placed under the control of the *S. pombe urg1* promoter and integrated with the Dam gene at the *urg1* locus (Fig. 1A). At low copper concentrations, Ace1 is inactive, limiting Dam expression to basal levels. Upon exposure to copper, Ace1 becomes activated, initiating Dam expression.

**Figure 1.**
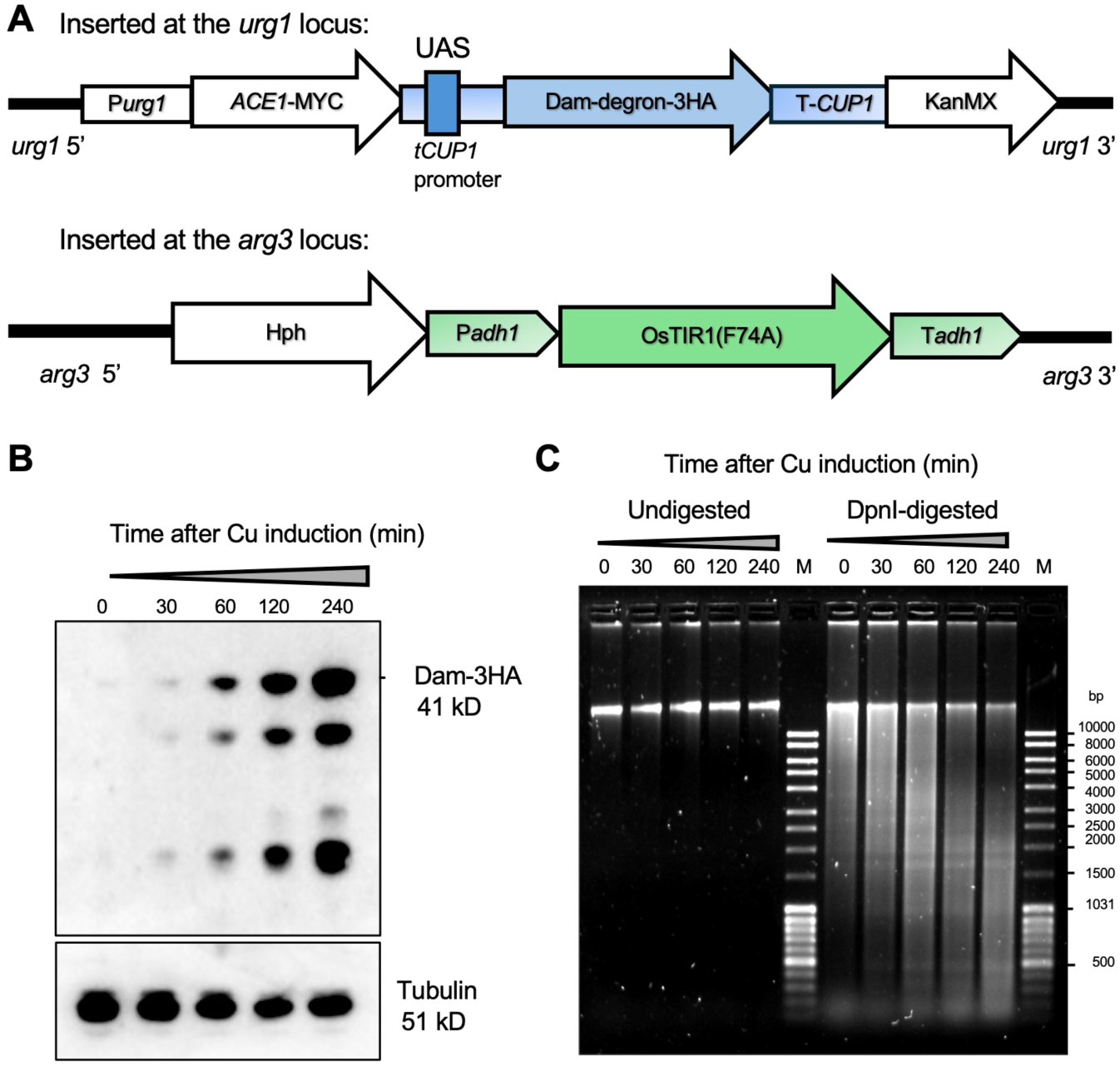
Copper induction of Dam expression in live *S. pombe* cells. (**A**) Integration cassettes. Upper: The copper-inducible Dam-degron-3HA integration cassette driven by the truncated *CUP1* (*tCUP1*) promoter with the *CUP1* terminator (T-*CUP1*). The *tCUP1* promoter contains a single upstream activating sequence (UAS) for copper-dependent binding by the Ace1 transcription factor also encoded in the cassette under the control of the *urg1* promoter. The KanMX gene allows G-418 selection for integration at the *urg1* locus. The Dam open reading frame is fused to an auxin-dependent degron to reduce basal expression in the presence of 5a-IAA, and three HA tags for detection. Lower: The *adh1* promoter drives constitutive OsTIR1(F74A) expression; TIR1(F74A)-induced degradation of Dam requires 5a-IAA. The Hph gene allows hygromycin selection for integration at the *arg3* locus. (**B**) Immunoblot analysis of Dam-degron-3HA expression at different time points following copper induction. (**C**) Agarose gel electrophoresis of genomic DNA. At left: genomic DNA before DpnI digestion; at right: DpnI-digested genomic DNA. The DNA is almost completely digested by DpnI after 240 min, indicating that it is almost fully methylated in vivo. M: DNA marker.

However, achieving strongly regulated Dam expression was compromised by high background GATC site methylation. To address this, we fused Dam to an auxin-inducible degron with three HA tags (3HA) (Fig. 1A). We integrated a gene encoding an OsTIR1 F-box protein with the F74A mutation in the auxin-binding motif at the *arg3* locus (Fig. 1A). This mutation enhances its specificity for a synthetic auxin, 5-adamantyl indole acetic acid (5a-IAA) over endogenous auxin, enabling conditional degradation of Dam in the presence of 5a-IAA (55). By employing this method, we achieved precise control over Dam protein expression, minimized background methylation and successfully detected the copper-dependent induction of Dam-degron-3HA with an anti-HA antibody (Fig. 1B).

To estimate the relative expression of Dam in *S. pombe* cells, we compared Dam-3HA levels with those of a native protein. We selected Gcn5, a histone acetyltransferase, which exhibits intermediate expression levels in exponentially growing cells (49,50). Both Dam and Gcn5 were tagged with the same 3HA epitope to ensure uniform detection, and their relative expression was analyzed by immunoblotting (Supplementary Fig. S1). Densitometric analysis revealed that Dam-3HA accumulated at 3.1-fold higher levels (replicate: 1.8-fold) compared to Gcn5-3HA. Given that Gcn5 itself is a moderately expressed protein, the observed Dam abundance is comparatively low relative to the bulk proteome of *S. pombe* (note the log_10_ scale). Various sequence-specific transcription factors are expressed at similar protein levels (Supplementary Fig. S1B).

### Euchromatin is generally accessible in living *S. pombe* cells

To assess genome accessibility, exponentially growing *S. pombe* cells were treated with 5a-IAA, followed by its removal and subsequent copper-induced expression of Dam. Time-course experiments were performed to monitor expression dynamics and methylation rate. Immunoblotting confirmed robust Dam induction (Fig. 1B), while DpnI digestion of genomic DNA, which specifically cuts only at GATC sites methylated on both strands, revealed progressively increasing methylation at successive time points. Almost complete digestion was observed after 240 min, indicating that most of the *S. pombe* genome is accessible to Dam in vivo (Fig. 1C).

This observation was strengthened by a comprehensive analysis of the 32,949 GATC sites in the *S. pombe* genome, utilizing quantitative DNA accessibility sequencing (qDA-seq) (33,34) (Fig. 2). Purified genomic DNA is subjected to DpnI digestion, followed by sonication to reduce DNA fragment size, and Illumina paired-end sequencing. We quantified the fraction methylated for every GATC site. At each time point, the fraction of DpnI-generated cuts was determined by counting the number of molecules beginning or ending within the GATC site and dividing by the total DNA fragment coverage at that site. The methylated fraction data for all GATC sites at each time point were combined to generate median plots for various genomic regions. The methylation time course for GATC sites in gene bodies is presented as an example (Fig. 2A): the median GATC site is > 70% methylated over the time course, although there is a wide range in methylation rate. The trend towards complete methylation is clear, but it was not attained presumably because not enough Dam was produced during the time course, since the methylation rate depends on the Dam concentration. To quantify the methylation rate for each genomic region, we plotted the natural log of the median unmethylated fraction against time, assuming pseudo-first order kinetics (validated by the straight lines observed; Fig. 2B). The slope of the regression line is a measure of the apparent rate constant, which was normalized to the genomic median rate (set at 1.0). GATC sites in promoters and tRNA genes are methylated faster than those in gene bodies, as expected, given that promoters and tRNA genes are often nucleosome-depleted. The median and range of rate constants for all individual GATC sites in each genomic region are presented as box plots (Fig. 2C).

**Figure 2.**
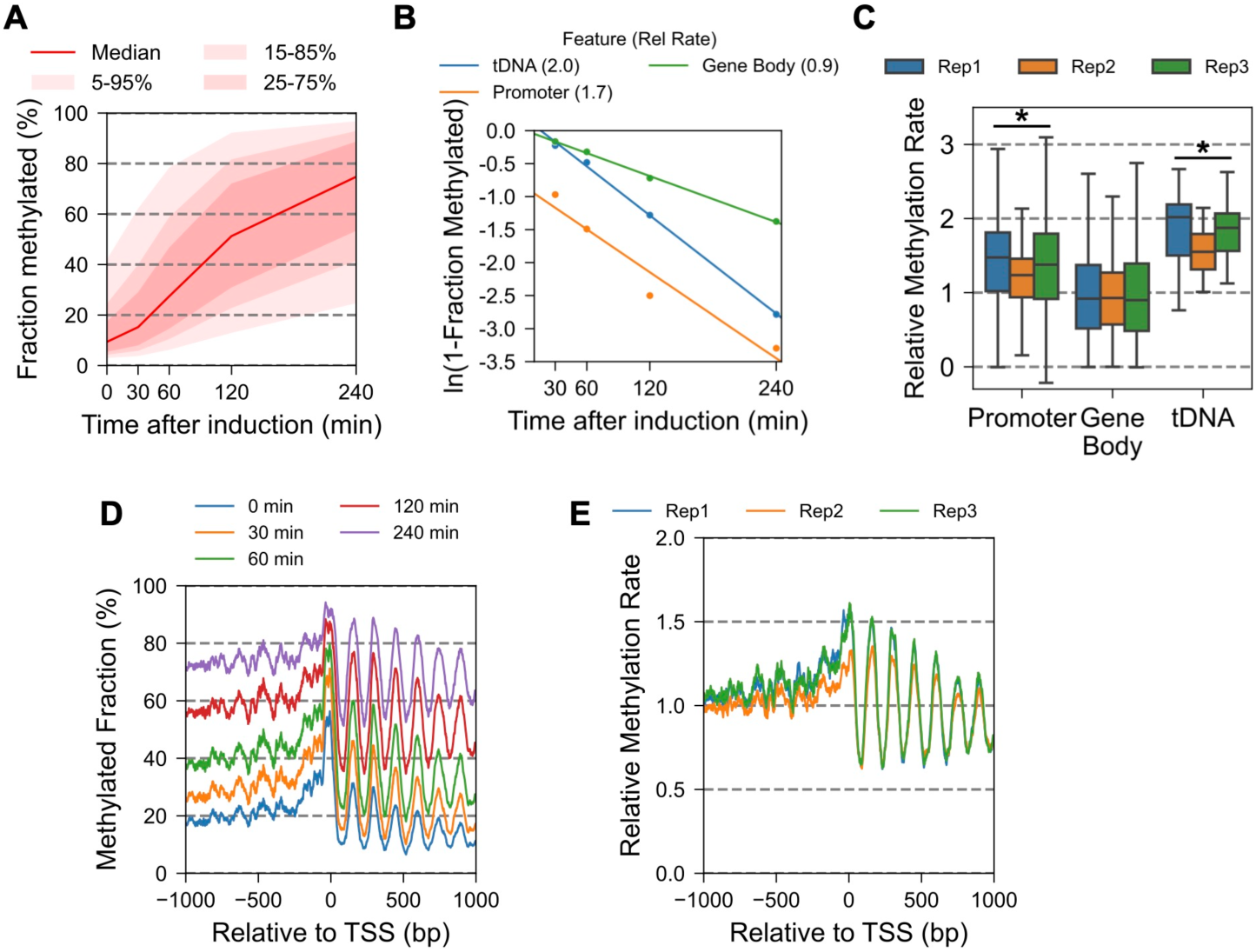
Euchromatin is generally accessible in live *S. pombe* cells. (**A**) Global analysis of Dam methylation of GATC sites located in gene bodies shows a trend towards complete methylation. Red line: methylation of the median GATC site; red shading: data range. (**B**) Dam methylation rate analysis of the median GATC site in promoters, gene bodies and tRNA genes. (**C**) Promoters and tRNA genes are methylated faster than gene bodies. Box plots showing the distribution of individual GATC site methylation rates for GATC sites in promoters (defined as 100 bp upstream from the TSS), gene bodies and tRNA genes for three biological replicate experiments. Each box contains 25 to 75% of the data, the central line is the median, and the whiskers represent 1.5 times the interquartile range to the farthest data points. Asterisk indicates that the feature is significantly different from gene bodies (Mann-Whitney U test). (**D**) Nucleosome phasing in vivo: Time course analysis of average Dam methylation at GATC sites for all genes aligned on the TSS, smoothed with a 21-bp window. (**E**) Nucleosome phasing in vivo: relative average methylation rate at GATC sites for all genes aligned on the TSS (smoothed with a 21-bp window) for three biological replicate experiments.

To analyze the chromatin organization of the 5,116 protein-coding genes in vivo, we plotted the mean GATC site methylation profile for each time point relative to the TSS of each gene. We observed a clear nucleosome phasing pattern downstream of the TSS at each time point (Fig. 2D). The peaks represent fast-methylating linkers; the troughs represent slower-methylating nucleosomal DNA. The profile shifts upwards with each time point as the genes become more and more methylated. Average methylation rates were normalized to the genomic median rate (set at 1), and three biological replicate experiments were compared (Fig. 2E). Estimation of the nucleosome spacing at ∼149 bp confirms that the nucleosomes are unusually close together on gene bodies in *S. pombe*, with very little linker. Estimates for nuclei range from 150 to 156 bp, derived from MNase data (30–32) and chemical mapping data (56). Overall, our results demonstrate that euchromatic regions (promoters, gene bodies and tRNA genes) are accessible to Dam methylase in live *S. pombe* cells.

To examine whether transcription influences genome accessibility, promoters and gene bodies were divided into quintiles based on their RNA-seq expression levels in a 972h-strain grown in a similar medium without added Cu (data from (51)). Quintile Q1 represents genes with the lowest expression, whereas Q5 contains genes with the highest expression. Box plots showing the distribution of methylation rates for the GATC sites in each gene quintile indicate that the median rate for gene bodies is unaffected by transcription level (Supplementary Fig. S2A). We conclude that transcriptional activity has little effect on methylation rate.

To assess whether DNA replication has an impact on genome accessibility, cells were arrested in uncommitted G1 by culturing in medium lacking nitrogen (57), followed by Dam induction and time course analysis (Supplementary Fig. S3). A high rate of methylation was also observed in arrested cells, indicating that DNA replication does not make a major contribution to global genome accessibility in vivo. We note that methylation rates in log phase and arrested cells should not be compared directly because the amount of Dam produced and the kinetics of Dam production may be different (34,37).

### Pericentromeric heterochromatin is accessible and methylated at a similar rate to euchromatin

The basic organization of *S. pombe* centromeres is illustrated in Fig. 3A. The central domain consists of the non-repetitive central core (*cnt*), which is assembled into both canonical nucleosomes and centromeric nucleosomes containing the centromere-specific H3 variant Cnp1 (58) (CENP-A in higher organisms), flanked by the innermost repeats (*imr*) which are, in turn, flanked by the pericentromeric *dg* and *dh* repetitive elements. Pericentromeric heterochromatin is composed of canonical nucleosomes marked by H3K9me2/3 and associated with *S. pombe* HP1 (Swi6). Based on our prior studies in budding yeast and human cells, centromeric chromatin was expected to be relatively inaccessible, characterized by slow methylation rates (16,34). However, *cnt* and *imr* display faster methylation (Fig. 3B), at ∼1.5 times the genomic median rate (Fig. 3C,D), indicating that *S. pombe* centromeric chromatin differs from that of budding yeast and human MCF cells. The heterochromatic *dg* repeats are methylated at about the same rate as gene bodies, whereas the heterochromatic *dh* repeats are methylated slightly faster than gene bodies (Fig. 3B-D). In conclusion, the centromeric central core and innermost repeats are methylated slightly faster than gene bodies, whereas the heterochromatic *dg* and *dh* repeats are methylated at rates comparable to gene bodies, indicating euchromatin-like accessibility.

**Figure 3.**
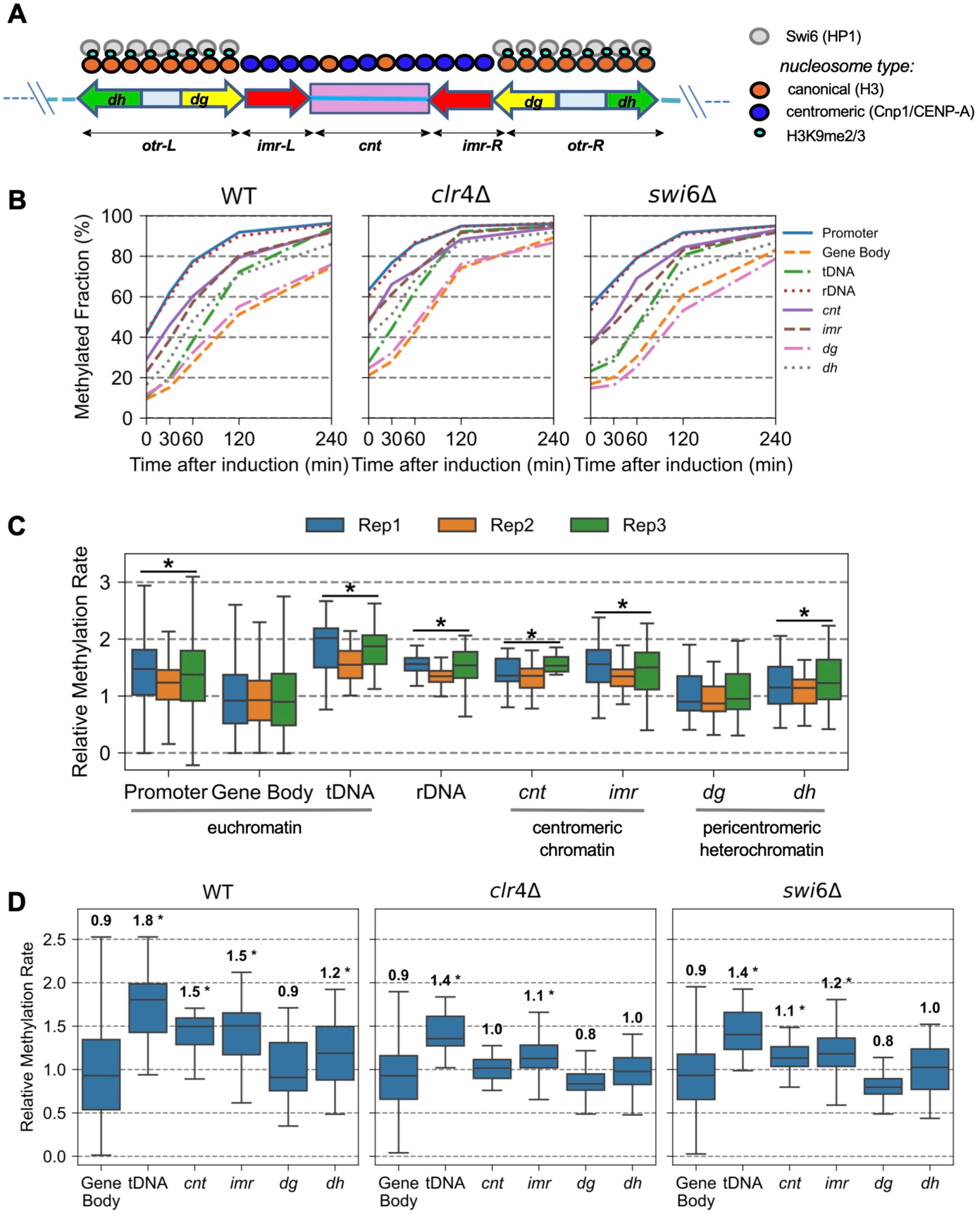
Pericentromeric heterochromatin is accessible in live *S. pombe* cells. (**A**) Schematic representation of *S. pombe* centromeres: the centromeric core consists of a unique central region (*cnt*) flanked by innermost repeats (*imr*), which are flanked in turn by pericentromeric heterochromatin (*otr:* outermost repeats) composed of *dg* and *dh* repeats. (**B**) Centromeres and pericentromeres are methylated at a similar rate or faster than gene bodies in wild type, *clr4Δ* and *swiCΔ* cells. Dam methylation time courses showing the median rate for various genomic regions in vivo. (**C**) Box plots showing the distribution of methylation rates for GATC sites in different genomic regions (the euchromatin data are the same as in Fig. 2C). Each box contains 25 to 75% of the data, the central line is the median, and the whiskers represent 1.5 times the interquartile range to the farthest data points. Asterisk indicates that the feature is significantly different from gene bodies (Mann-Whitney U test). (**D**) Average relative methylation rates in WT, *clr4Δ* and *swiCΔ* cells. Boxes indicate average methylation rates for three biological replicate experiments (see Supplementary Fig. S4 for a comparison of the biological replicate experiments). Numbers indicate the median value. Boxes and asterisks as in *C*.

Chromosome III carries 100-150 copies of rDNA at both ends. Each copy contains a gene which is transcribed by RNA polymerase I (Pol I) and processed to produce the 18S, 5.8S and 28S rRNAs. The rDNA loci are marked by H3K4me and weakly enriched with H3K9me (22,59), probably reflecting the presence of both active nucleosome-depleted rDNA repeats (H3K4me) and inactive fully nucleosomal rDNA repeats (H3K9me), as is the case in budding yeast (60). The rDNA repeats are methylated relatively fast (Fig. 3B,C). This observation most likely reflects the weighted average of the median methylation rates of active and inactive rDNA repeats.

### Loss of Clr4 or Swi6 does not increase heterochromatin accessibility in vivo

Our observation that heterochromatin and euchromatin are methylated at similar rates in vivo implies that Clr4-mediated H3K9 methylation and consequent HP1/Swi6 binding have little effect on DNA accessibility in heterochromatin. To test this prediction, we repeated the experiment in *clr4* and *swi6* null mutants (Fig. 3B,D). All genomic regions are > 80% methylated at the end of the time course in both mutants (Fig. 3B), indicating that the *S. pombe* genome is globally accessible in vivo in the absence of Clr4 or Swi6.

Methylation rates in mutant strains should not be compared directly with wild type because they depend on the amount of Dam produced and the induction kinetics, which may be different (16,34,37). However, we can compare relative rates for different genomic regions after normalisation to the genomic median rate within each data set (Fig. 3D). The box plots show the average rate for each GATC site relative to the genomic median from three biological replicate experiments for wild type (left panel), *clr4Δ* (middle panel) and *swiCΔ* cells (right panel). The individual replicates are compared in Supplementary Fig. S4. We omitted the data for promoters and rDNA in Fig. 3D, because methylation of these regions reaches a limit at ∼95% after 120 min (Fig. 3B), resulting in relatively poor regression lines for the rate measurements. This is because the methylation level prior to induction is higher in both mutants than in wild type (Fig. 3B). The reason for the high background is unclear; higher 5a-IAA concentrations did not suppress the background methylation in *clr4Δ* cells (Supplementary Fig. S5A,B).

Relative methylation rates for the various genomic regions in wild type, *clr4Δ* and *swiCΔ* cells show similar trends (Fig. 3D). In wild type cells, GATC sites in the heterochromatic *dg* elements are methylated at about the same rate as those in gene bodies (∼0.9 times the genomic median rate), whereas sites in *dh* elements are methylated a little faster than those in gene bodies (∼1.2 times the genomic median rate) (Fig. 3D, left panel). In *clr4Δ* cells, *dg* elements (∼0.8x) and *dh* elements (∼1.0x) are methylated at about the same rate as gene bodies (∼0.9x) (Fig. 3D, middle panel). The same is true in *swiCΔ* cells (right panel): *dg* (∼0.8x), *dh* (∼1.0x) and gene bodies (∼0.9x). Thus, loss of Clr4 or Swi6 results in a slightly decreased methylation rate at *dh* elements and no change in *dg* element methylation rate, relative to gene bodies. Since faster methylation is predicted if Clr4 and Swi6 reduce DNA accessibility in heterochromatin, our data indicate that Clr4 and Swi6 do not limit DNA accessibility in heterochromatin in vivo.

Interestingly, genomic regions lacking Clr4 and Swi6 are also affected in the mutants (Fig. 3D). In wild type cells, tRNA genes are methylated at ∼1.8 times the genomic median (gene bodies: ∼0.9x), whereas in *clr4Δ* and *swiCΔ* cells, they are methylated more slowly, at ∼1.4 times the genomic median (gene bodies: ∼0.9x). Similarly, *cnt* is methylated at ∼1.0 times the genomic median in *clr4Δ* cells and ∼1.1 times the genomic median in *swiCΔ* cells, whereas *cnt* is methylated at ∼1.5 times the genomic median in wild type cells. Methylation rates at *imr* sites show a similar trend.

In summary, relative to the genomic median rate, heterochromatin (*dg* and *dh*) and centromeres (*cnt* and *imr*) are methylated slightly slower in the mutants than in wild type. Most importantly, we conclude that Clr4 or Swi6 do not limit heterochromatin accessibility in vivo.

### The mating type loci are methylated at similar rates to euchromatin

The silent mating type locus comprises a ∼20 kb region flanked by inverted repeat (IR) sequences, incorporated into canonical nucleosomes marked by H3K9me2/3 and bound by HP1 (Swi6) (21,61,62). It includes the *mat2-P*, *mat3-M* and *cen-H* loci (Fig. 4A). The *cen-H* locus shares extensive homology with the pericentromeric *dg* and *dh* elements (62). The euchromatic active *mat1* locus is located upstream of the silenced locus on chromosome II. Since the wild type strain (972h-) used in the experiments above lacks the mating type loci, we used a *mat1-Msmt0* strain (mating type M) instead. This strain contains the *mat3-M* sequence at the active *mat-1* locus and an identical silenced copy at the *mat3-M* locus (63). The *mat2-P* locus is a silenced copy of the mating type P sequence.

**Figure 4.**
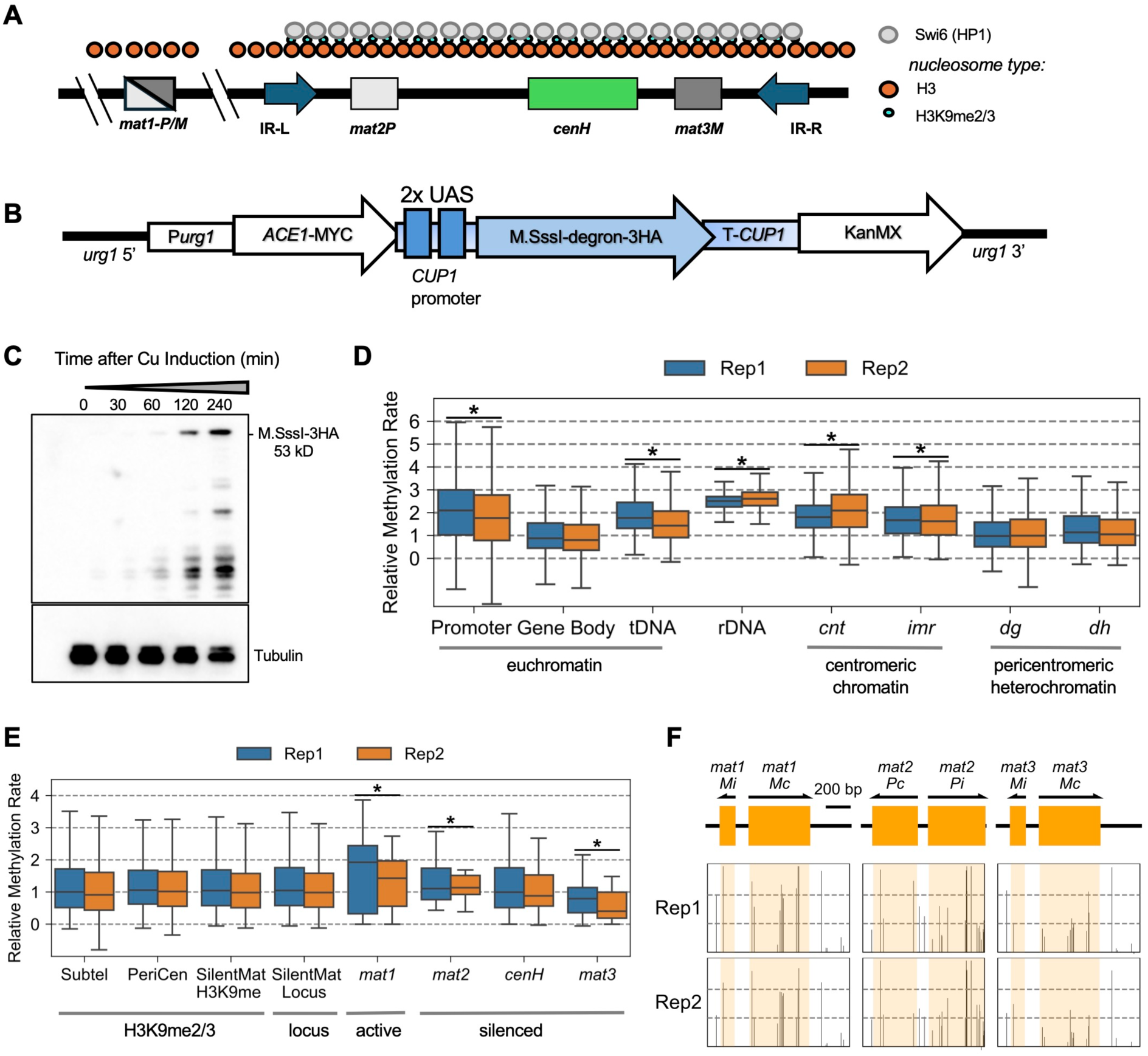
Mating type locus (*mat*) heterochromatin is accessible in live *S. pombe* cells. (**A**) Heterochromatin organization at the mating type locus, comprising the transcriptionally active *mat1M* region and the ∼20 kb domain containing the silent *mat2P*, *cenH* and *mat3M* loci bordered by inverted repeats. This silent domain is enriched in H3K9me2/3 and bound by Swi6. (**B**) Schematic of the inducible M.SssI-degron-3HA integration cassette, driven by the full-length *CUP1* promoter, which has two UAS elements. (**C**) Immunoblot showing Cu-induced expression of M.SssI-degron-3HA. (**D**) Centromeres and pericentromeres are methylated by M.SssI at a similar rate or faster than gene bodies. Box plots showing the distribution of methylation rates for CG sites in various genomic regions (two biological replicate experiments). Each box contains 25 to 75% of the data, the central line is the median, and the whiskers represent 1.5 times the interquartile range to the farthest data points. A few CG sites have regression lines with poor correlations resulting in negative rates. Asterisk indicates that the feature is significantly different from gene bodies (Mann-Whitney U test). (**E**) The silent mating type locus is methylated at a similar rate to gene bodies, except for the *mat3M* locus. Box plots showing the distribution of methylation rates for CG sites in the mating type locus. Box plots for sub-telomeric (Subtel), pericentromeric (PeriCen) and the silent mating type locus (SilentMat) regions defined by H3K9me2/3 ChIP-seq data (43). The mating type loci are defined by their genomic annotations. (**F**) M.SssI methylation rates for the active *mat1M* and the silenced *mat2P* and *mat3M* loci, showing the locations of the silenced genes. Each CG site in the locus is shown as a column indicating the relative rate at that site (lower dashed line: genomic median set at 1).

Since the *mat2-P*, *mat3-M* and *cen-H* loci have a low density of GATC sites, we elected to use M.SssI instead of Dam. M.SssI methylates C in CG dinucleotides to 5-methylcytosine (5mC), enabling higher resolution mapping, because CG motifs are much more common than GATC motifs. We used Nanopore long-read sequencing because it can detect 5mCG and it can distinguish between the identical *mat1-M* and *mat3-M* loci. Initial experiments using the truncated *CUP1* promoter to induce M.SssI resulted in low genomic methylation levels. To increase the amount of M.SssI produced, the truncated *CUP1* promoter was replaced with its full-length counterpart, which has two UASs recognized by Ace1 (Fig. 4B). M.SssI activity increased to sufficiently high levels (Fig. 4C,D), although it was still not as active as Dam.

The relative methylation rates for M.SssI and Dam at the various genomic regions are similar, but there are some differences (Fig. 4D; cf. Fig. 3C). In both cases, promoters, tRNA genes and rDNA are methylated faster than gene bodies, but at different relative rates for Dam and M.SssI. This is probably because M.SssI has many more target sites (CG) than Dam (GATC), resulting in more data for each genomic region (e.g. there are relatively few GATC sites in tRNA genes because they are so short). However, M.SssI activity levels in vivo are much lower than those of Dam, presumably due to differences in enzyme concentration, enzyme turnover number and substrate concentration.

Centromeric regions (*cnt* and *imr*) are methylated faster by M.SssI than gene bodies, whereas the pericentromeric heterochromatin (*dg* and *dh* repeats) is methylated at about the same rate as gene bodies (Fig. 4D). Analysis of the methylation rate distributions in the transcription quintiles shows that methylation rates in gene bodies are independent of the transcription level (Supplementary Fig. S2B). The higher resolution afforded by M.SssI data allowed us to construct a nucleosome phasing plot for each quintile, revealing that the least expressed genes (Q1) display weaker phasing than the genes in the other quintiles (Supplementary Fig. S2C). The reduced phasing in Q1 can, in principle, be explained by the absence from some of the genes of the putative barrier complex required to form an NDR, adjacent to which the nucleosomal array forms (64).

We quantified CG methylation rates in the mating type locus (Fig. 4E). The entire silenced locus (∼20 kb), as defined by the genomic sequence, is methylated at about the same rate as the rest of the genome, indicating that this heterochromatic region is accessible, just like pericentromeric heterochromatin. A similar result is obtained if the silenced mating type locus, pericentromeres and sub-telomeric regions are defined by the H3K9me2/3 mark (Fig. 4E). The silent *mat2-P* and *cenH* loci are methylated at similar rates to the entire silenced locus. However, the silent *mat3-M* locus is methylated somewhat more slowly than the rest of the locus, whereas the identical, active, *mat1-M* locus (located outside the silenced region) is methylated faster than the silenced region (Fig. 4E,F). This difference between the identical *mat3-M* and *mat1-M* sequences is primarily due to fast methylation within the coding regions (*Mi* and *Mc*) in *mat1-M* relative to *mat3-M*; the CG sites flanking *Mi* and *Mc* are methylated at about the same rate (Fig. 4F). The explanation for the difference between *mat3-M* and *mat1-M* is unclear; the active *mat1-M* locus might be nucleosome-depleted relative to *mat3-M*. We conclude that the mating type loci are generally accessible, even though subregions are methylated at different rates.

### Condensed knob chromatin is accessible in vivo

Condensed chromatin regions lacking H3K9me and Swi6 (termed “knobs”) are located adjacent to the sub-telomeres of chromosomes I and II (29). Knob regions span ∼50 kb. Their telomere-proximal boundaries are defined by the absence of sub-telomeric H3K9me3, but the telomere-distal boundaries are unclear. Consequently, we divided the 50-kb region proximal to each telomere into 10-kb windows (termed K1 to K5) and made box plots of CG site methylation rates within each window (Supplementary Fig. S6). The sub-telomeric region of each of the four telomeres is methylated at rates close to the genomic median rate. At the left telomere of chromosome I, the K2 and K3 windows are methylated slightly faster than the genomic median rate. All five windows at the right telomere of chromosome I and at the left telomere of chromosome II are methylated close to the genomic median rate. At the right telomere of chromosome II, the K2 window is methylated slightly more slowly than the genomic median rate. In summary, the observed methylation rates do not suggest that condensed knob chromatin is less accessible in vivo.

### The genome is mostly inaccessible in wild type, *clr4Δ* and *swiCΔ* nuclei

We measured genome accessibility in nuclei isolated from wild type, *clr4Δ* and *swiCΔ* cells. Nuclei were incubated with purified Dam at various concentrations. Genomic DNA was extracted and digested with DpnI as in live cell experiments. Dam-treated DNA from wild type nuclei was only partially digested by DpnI, indicating that a substantial fraction of the DNA was unmethylated, even at the highest Dam concentration (Fig. 5A).

**Figure 5.**
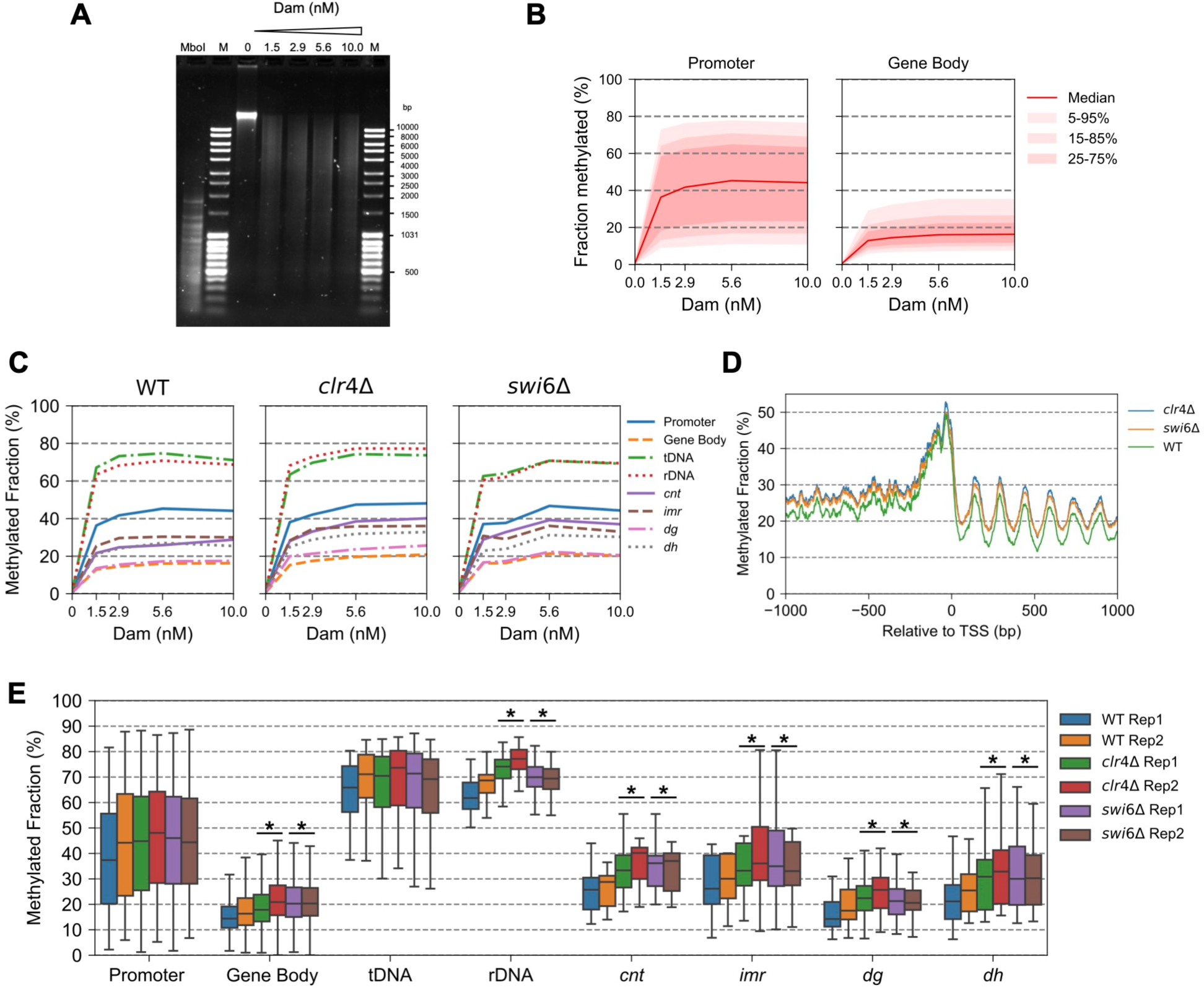
The genome is mostly inaccessible in wild type, *clr4Δ* and *swiCΔ* nuclei. Dam methylation of genomic DNA in nuclei reaches a limit. (**A**) Isolated wild type nuclei were treated with increasing amounts of Dam. Purified genomic DNA was digested with DpnI and resolved in a 1% agarose gel stained with SYBR Gold. MboI: marker for complete digestion of GATC sites (purified unmethylated genomic DNA was fully digested with MboI). M: marker. (**B**) Wild type nuclei: Global analysis of Dam methylation of GATC sites located in promoters and gene bodies. Red line: methylation of the median GATC site; red shading: data range. (**C**) Median methylation of various genomic regions as a function of Dam concentration for wild type, *clr4Δ* and *swiCΔ* nuclei. (**D**) Nucleosome phasing detected by Dam methylation of nuclei. Average methylation at the highest Dam concentration in wild type, *clr4Δ* and *swiCΔ* nuclei for all genes aligned on the TSS, smoothed with a 21-bp window. (**E**) Median methylated fractions of various genomic regions at the highest Dam concentration in wild type, *clr4Δ* and *swiCΔ* nuclei (data for two biological replicates). Asterisk indicates that the feature in the mutant is significantly different from that in wild type (Mann-Whitney U test).

Genome-wide analysis confirmed that methylation is limited in wild type nuclei, unlike in vivo. Increasing amounts of Dam do not result in increased methylation in nuclei, consistent with protection of nucleosomal DNA from Dam (Fig. 5B). Median methylation at GATC sites reached a plateau at 44% in promoters and 16% in gene bodies. These numbers indicate that the median promoter GATC site is accessible in 44% of nuclei and inaccessible in 56% of nuclei. Similarly, the median gene body GATC site is accessible in 16% of nuclei and inaccessible in 84% of nuclei. Promoter methylation is higher than in gene bodies because of the NDR present at most promoters. Gene body methylation is very low in nuclei presumably because there is very little linker DNA between nucleosomes on gene bodies (30–32,56). These observations indicate that a large fraction of the genome is completely inaccessible in isolated nuclei, unlike in vivo.

The median methylated fraction of promoters in *clr4Δ* and *swiCΔ* nuclei is similar to that of wild type nuclei, whereas that of gene bodies is slightly higher in the mutants than in wild type nuclei (Fig. 5C; Supplementary Fig. S5C). Nucleosome phasing plots for all genes at the highest Dam concentration show that the promoter NDR reaches a limit at ∼50% accessible in wild type, *clr4Δ* and *swiCΔ* nuclei, indicating that promoter GATC sites are inaccessible in about half of the nuclei (Fig. 5D). Nucleosome spacing is very similar in wild type and the mutants, but the mutants have a higher accessible fraction, suggesting that nucleosome occupancy is somewhat lower in *clr4Δ* and *swiCΔ* nuclei.

The tRNA genes and the rDNA are the most accessible regions in nuclei from wild type and both mutants, reaching a limit methylation of ∼70% (Fig. 5C,E). This very high level of methylation also indicates that at least 70% of the nuclei were exposed to Dam, and that the very low level of gene body methylation is not due to a problem with Dam entering the nuclei. The tRNA genes may be mostly accessible in nuclei because they are highly transcribed by RNA polymerase III and, at least in budding yeast, are occupied by the TFIIIB and TFIIIC transcription factors instead of a nucleosome (65), which may provide less protection from methylation. The high accessibility of tRNA genes in nuclei is consistent with their rapid methylation in vivo (Fig. 3D).

The pericentromeric *dg/dh* repeats and the centromeric *cnt* and *imr* regions are mostly inaccessible in nuclei (Fig. 5C). The heterochromatic *dg* elements are slightly more accessible in the mutants (26% in *clr4Δ* and 21% in *swiCΔ*) than in wild type nuclei (18% methylated) (Fig. 5E). The heterochromatic *dh* elements are also slightly more accessible in the mutants (33% in *clr4Δ*, 30% in *swiCΔ* and 26% in wild type). Thus, in isolated nuclei, loss of Clr4 or Swi6 results in a small increase in the accessible fraction, consistent with a small amount of extra stable protection of heterochromatic DNA by these proteins. However, similar increases in the accessible fraction are also observed at non-heterochromatic regions (*cnt*, *imr*, gene bodies and the rDNA; Fig. 5E). These data suggest that Clr4 and Swi6 have a modest protective role in nuclei at the global level. This role is presumably at least partly indirect, since non-heterochromatic regions are affected.

## Discussion

In this study, we have established a strong inducible system in *S. pombe* that integrates copper-inducible transcriptional control with auxin-mediated protein degradation to reduce background expression. We co-expressed a mutant OsTIR1 (F74A) that responds selectively to the synthetic auxin 5a-IAA. This dual layer strategy adds a post-translational level of regulation designed to eliminate low levels of protein produced before induction. However, background expression is not completely suppressed. This novel inducible system is likely to be useful to the *S. pombe* community.

We measured DNA accessibility in *S. pombe* genome-wide and found that euchromatic regions are globally accessible in vivo on a time scale of minutes, similar to our observations in budding yeast and human cells (16,34). These findings align with our nucleosome dynamics model, which proposes a continuous nucleosome flux in vivo, resulting from nucleosome eviction and replacement, nucleosome sliding along DNA and/or nucleosome conformational changes (16,34). These are all known catalytic activities of ATP-dependent chromatin remodelers. We propose that remodeling transiently exposes the DNA motif (GATC or CG), thereby allowing methylation even within nucleosomes. We argue that in nuclei, the nucleosomes are static because ATP and other critical co-factors are lost. Consequently, only linker DNA remains accessible for methylation in nuclei, while nucleosomal DNA is protected.

We found that pericentromeric heterochromatin and the heterochromatic mating type loci are fully accessible in living *S. pombe* cells, unlike in isolated nuclei, where it is mostly inaccessible. Loss of Clr4 or Swi6, which are required for heterochromatin formation, has only weak effects on heterochromatin accessibility. In isolated nuclei, loss of Clr4 or Swi6 results in a small increase in the accessible fraction, possibly explained by loss of HP1/Swi6 dynamics at heterochromatic regions, and by a small decrease in nucleosome occupancy elsewhere. These effects must be at least partly indirect since they extend to regions where Clr4 and Swi6 do not bind.

Loss of Clr4 or Swi6 in vivo results in somewhat slower methylation rates in most regions relative to gene bodies, implying that Clr4 and Swi6 facilitate access to DNA, rather than prevent it. However, these rates are relative to gene bodies and the nuclei data suggest that gene bodies have lower nucleosome occupancy in both mutants. Lower nucleosome occupancy might be expected to increase the methylation rate in vivo, although such an effect might cancel out because we are normalising to the genomic median rate in vivo. An alternative explanation is based on the fact that nucleosomes are static in isolated nuclei but dynamic in vivo: nucleosome dynamics may be slower in *clr4Δ* and *swiCΔ* cells than in wild type cells. Clr4 and Swi6 might exert global effects on chromatin dynamics in vivo indirectly through 3-dimensional genome architecture, since heterochromatin loss in *clr4Δ* cells relaxes constraints within *S. pombe* chromosomes beyond heterochromatic regions (25). Overall, though, the effects of Clr4 or Swi6 loss on accessibility are quantitatively minor. The main conclusion is that H3K9 methylation and HP1/Swi6 do not limit DNA accessibility in living *S. pombe* cells.

Like *S. pombe* heterochromatin, human heterochromatin is accessible in vivo but, unlike *S. pombe* heterochromatin, it is methylated more slowly than euchromatin (16). This may reflect the fact that mammalian heterochromatin is condensed, unlike *S. pombe* heterochromatin (29). This observation implies that H3K9 methylation and HP1 do not by themselves cause heterochromatin condensation, which presumably requires additional factors, such as histone H1, which *S. pombe* lacks. HP1/Swi6 is quite mobile in *S. pombe* and murine cells, suggesting that there are windows of opportunity for accessing heterochromatic DNA (9,10,28). Our observations are clearly inconsistent with the hypothesis that heterochromatin is inaccessible in vivo.

The central role of RNAi in silencing shows that *S. pombe* heterochromatin must be transcribed at low levels, implying that it is accessible to the transcription machinery at least some of the time (24). Silencing is associated with reduced histone turnover in heterochromatin relative to euchromatin (24,66). Histone turnover or exchange is equivalent to nucleosome eviction and replacement in our flux model. Therefore, we might expect to observe reduced methylation rates at silenced loci. However, the methylation rates for gene bodies and heterochromatic regions are about the same in vivo, suggesting similar nucleosome dynamics. We suggest that remodeling activities involving nucleosome sliding and/or conformational changes are dominant over histone turnover at the silenced loci, resulting in little difference in methylation rate.

Heterochromatin in *S. pombe* differs from silenced chromatin in *S. cerevisiae*, which lacks HP1, H3K9me and RNAi. Instead, silencing in *S. cerevisiae* requires the Sir proteins, which *S. pombe* lacks. We have recently shown that the silenced regions in *S. cerevisiae* are methylated relatively slowly in wild type cells, and that loss of Sir2, Sir3 or Sir4 results in methylation rates similar to the rest of the genome, indicating that the Sir proteins impede, but do not prevent, access to silenced DNA (67). In contrast, here we show that *S. pombe* heterochromatin is methylated at similar rates to euchromatin in wild type cells.

In nuclei, most of the *S. pombe* genome is inaccessible, unlike in living cells. The very low accessibility of gene bodies is explained by the extremely short linker between nucleosomes, such that almost all of the DNA is protected by nucleosomes. In fact, the linker DNA is nominally only 2-6 bp, assuming a nucleosome core of 145-147 bp (68) and a spacing of 149-151 bp. If the average linker is really only 2-6 bp, steric clashes between adjacent nucleosomes are likely. Moreover, it is unlikely that the DNA methylases could access such short linkers. The limit methylation for gene bodies is 14 to 16%, indicating that ∼23 bp can be methylated per nucleosome if the spacing is 150 bp. Therefore, we propose that the terminal ∼10 bp on each side of the nucleosome tend to pull away from the histone octamer, effectively increasing the linker length and preventing steric clashes. In vitro, the terminal 10 bp on each side of the nucleosome undergo rapid cycles of association and dissociation (69,70); this effect may be magnified in *S. pombe* chromatin.

Analysis of our data for nuclei from different cell types reveals that the methylation limit value is related to the linker length (Fig. 6). We measured the average nucleosome spacing by fitting a decaying sine wave (37) to our Dam nucleosome phasing data for *S. pombe* genes (149 bp; replicate: 151 bp), *S. cerevisiae* genes (168 bp for two replicates) (34) and active genes in human MCF7 cells (184 and 185 bp) (16). The limit methylated fraction is given by [accessible DNA/(accessible + inaccessible DNA)]; the denominator is the nucleosome spacing. A plot of the observed limit methylated fraction against (1/nucleosome spacing) gives a straight line, indicating that the limit methylated fraction increases with linker length. The inaccessible fraction is derived from the reciprocal of the x-axis intercept (i.e., when the limit methylated fraction = 0), giving a value of ∼138 bp. This value is somewhat smaller than the nucleosome (∼147 bp) and is consistent with some methylation of nucleosomal terminal DNA due to breathing, in addition to the linker in nuclei from all three cell types. Thus, the gene body methylation limit in nuclei is determined by the linker length.

**Figure 6.**
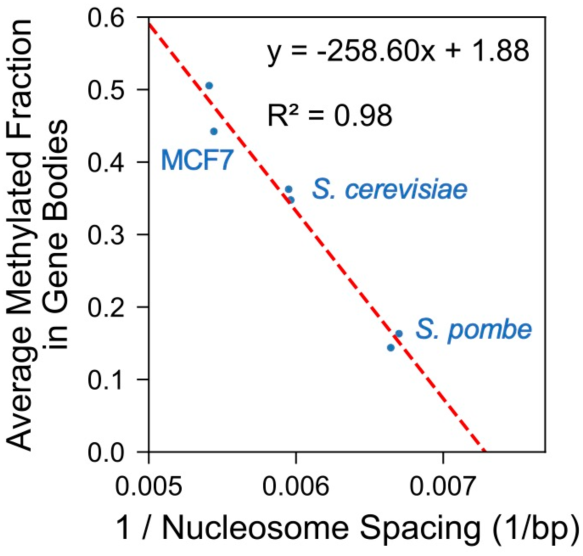
Linker DNA length determines the methylation limit in nuclei isolated from budding yeast, fission yeast and human MCF7 cells. Plot of the average methylated fraction in nuclei for *S. pombe* genes, *S. cerevisiae* genes and active genes in human MCF7 cells against the reciprocal of the average nucleosome spacing in nuclei. Data for two biological replicate experiments are shown for each cell type.

The accessibility of centromeric chromatin varies among species. In human MCF7 and MCF10A cells, centromeres are accessible, but methylated slower than gene bodies. In budding yeast, the centromeres are completely protected in vivo, although they are atypical, being point centromeres of ∼125 bp occupied by a single centromeric nucleosome with no linker included (71). By contrast, in *S. pombe*, the centromeric cores span 35 - 110 kb and are methylated faster than gene bodies. This increased methylation rate might reflect a global effect of heterochromatin on 3-dimensional genome architecture (25).

## Acknowledgements

We thank Shiv Grewal for strain SPT999a. We thank Hemant Prajapati, Peter Eriksson and Kenneth Wu for advice and comments. This study used the high-performance computational capabilities of the Biowulf Linux cluster at the NIH.

## Author contributions

Conceptualization - D.C.; Investigation - L.P.; Formal analysis and software - Z.X.; Writing - original draft - L.P.; Writing - review and editing - L.P. and D.C.

## Supplementary data

Supplementary Figs. S1 - S6 and Table S1.

## Data availability

The Illumina and Nanopore sequence data generated in this study have been deposited in the GEO database with accession code GSE313347.

The code used to analyse the data is available at FigShare.

