## Supplemental Information for "Histone H3K9 methylation and Heterochromatin Protein 1 do not limit DNA accessibility in living *S. pombe* cells"

**Lalita Panigrahi[1], Zhuwei Xu[1] and David J. Clark***
Division of Developmental Biology, *Eunice Kennedy-Shriver* National Institute of Child Health and Human Development, National Institutes of Health, Bethesda MD 20892, USA

[1] These authors contributed equally.

### Supplementary Figures S1-S6.

### Supplementary Table S1.

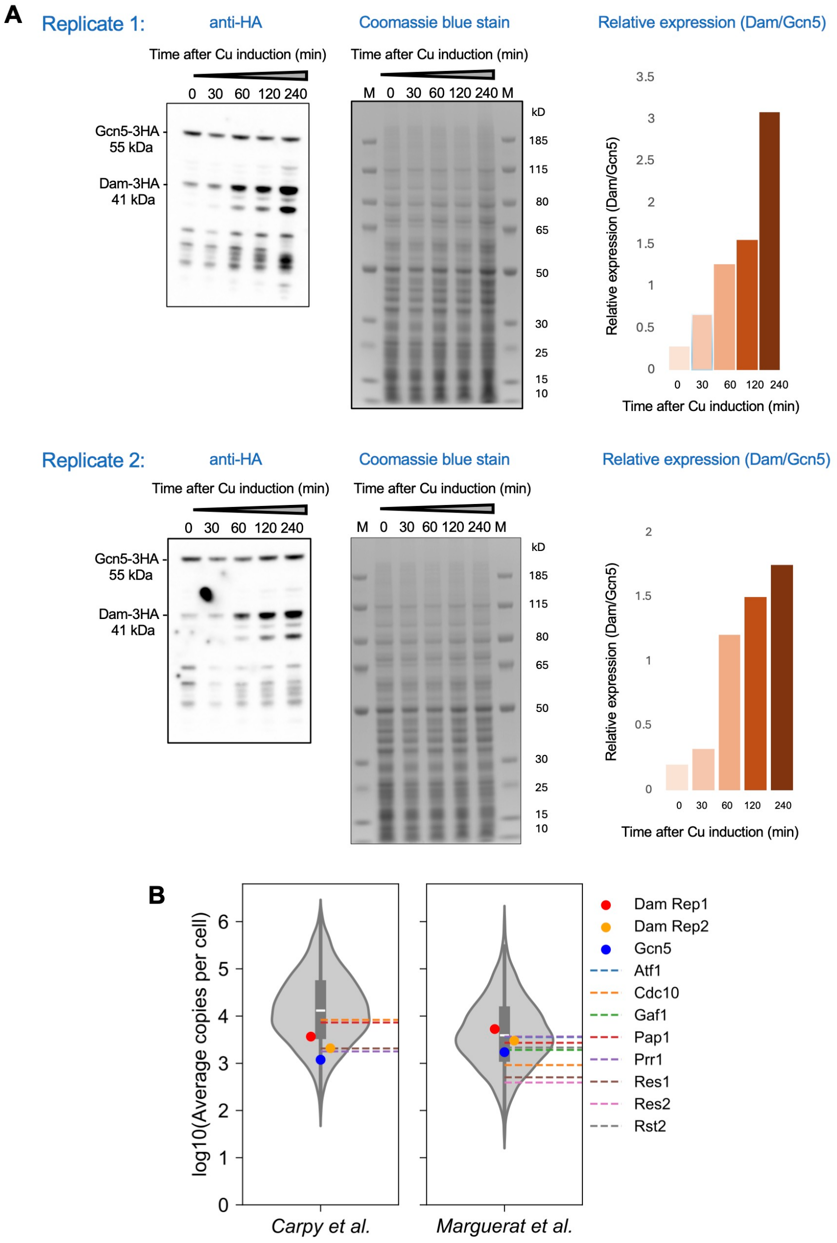


**Supplementary Fig. S1.** Dam-HA is expressed at similar levels to Gcn5 in *S. pombe*. (**A**) Immunoblot analysis of Dam-3HA and Gcn5-3HA (internal control) at different time points after induction (left panels). Coomassie-stained SDS-PAGE gel of the same extract used for immunoblotting, shown as a loading control, to verify equal protein amounts in each lane (middle panels). Densitometric analysis of the immunoblots revealed that Dam-3HA accumulates to ~2 to 3-fold higher levels than Gcn5-3HA (right panels). (**B**) Violin plots showing the distribution of *S. pombe* protein abundance in cells in vegetative growth (data from (49,50) obtained from Pombase), with the amount of Dam after 240 min induction relative to Gcn5 superimposed. Note the log10 scale. After normalizing Dam to Gcn5, the estimated Dam abundance falls within the lower to intermediate region of the protein distribution range. Data for the two biological replicates above are shown (orange and red for Dam; blue for Gcn5). For comparison, protein expression data for eight sequence-specific transcription factors are shown (dashed lines).

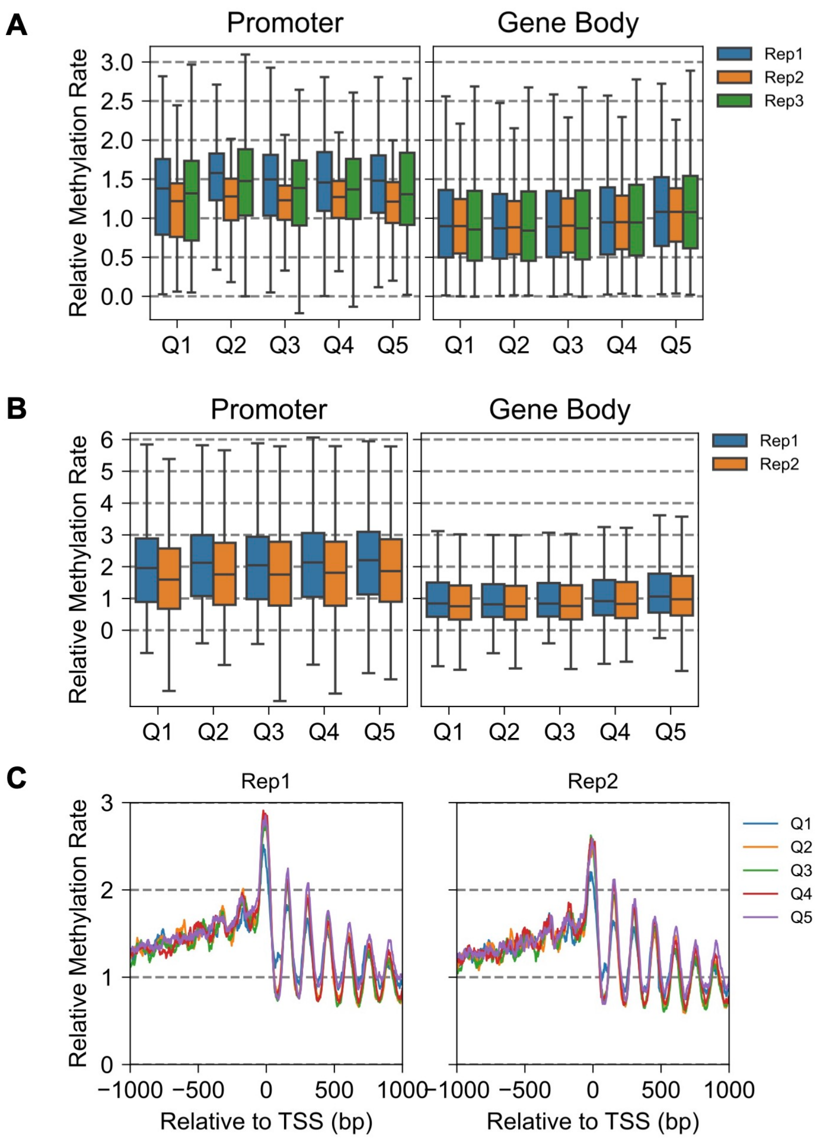


**Supplementary Fig. S2.** No significant effect of transcription level on methylation rates in live *S. pombe* cells. Protein coding genes were divided into quintiles according to their transcription level (RNA-seq data from (51)). The RNA-seq data were obtained for a 972h- strain grown in a similar medium (0.5% yeast extract, 3% glucose with adenine, histidine, leucine, uracil and lysine) without added Cu (51). Q5 contains the most highly transcribed genes. (**A**) Dam data in box plots for promoters and gene bodies. Each box contains 25 to 75% of the data, the central line is the median, and the whiskers represent 1.5 times the interquartile range to the farthest data points. Quintiles Q2 - Q5 are not statistically different from Q1 (Mann-Whitney U test). (**B**) M.SssI data in box plots for promoters and gene bodies. Quintiles Q2 - Q5 are not statistically different from Q1 (Mann-Whitney U test). (**C**) Relative M.SssI methylation rates for the genes in each quintile aligned on the TSS as a function of distance from the TSS, smoothed with a 21-bp window.

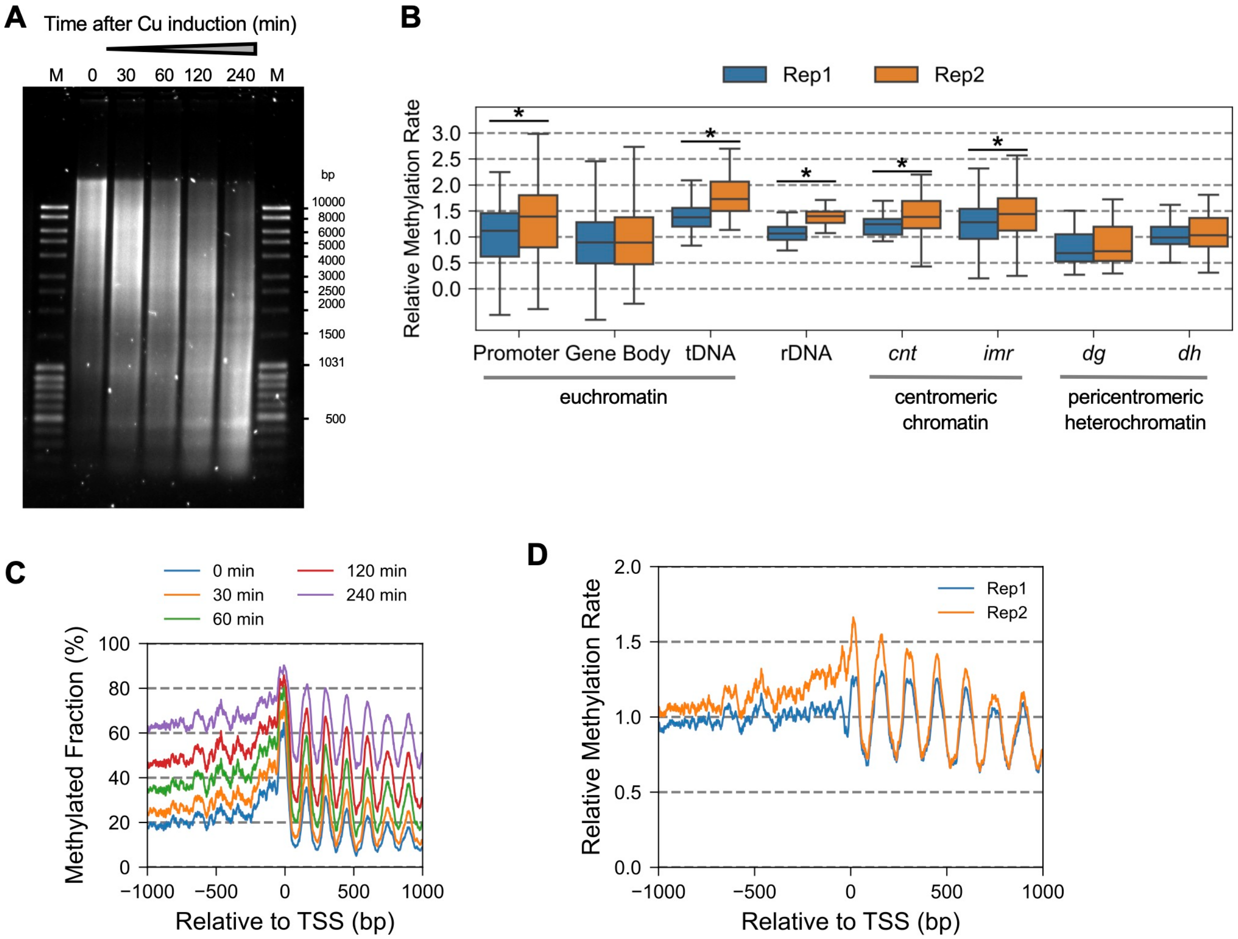


**Supplementary Fig. S3.** The *S. pombe* genome is globally accessible in arrested wild type cells.
(**A**) Genomic DNA methylation after Dam induction in cells arrested by nitrogen starvation. Purified genomic DNA was digested with DpnI. Samples were resolved on 1% agarose gel visualized using SYBR Gold staining. M: DNA marker. (**B**) Box plots showing the distribution of GATC site methylation rates in various genomic regions. Each box contains 25 to 75% of the data, the central line is the median, and the whiskers represent 1.5 times the interquartile range to the farthest data points. Asterisk indicates that the feature is significantly different from gene bodies (Mann-Whitney U test).
(**C**) Nucleosome phasing in arrested cells (replicate 1): Time course analysis of average Dam methylation at GATC sites for all genes aligned on the TSS, smoothed with a 21-bp window.
(**D**) Nucleosome phasing in arrested cells: relative average methylation rate at GATC sites for all genes aligned on the TSS (smoothed with a 21-bp window) for two biological replicate experiments.

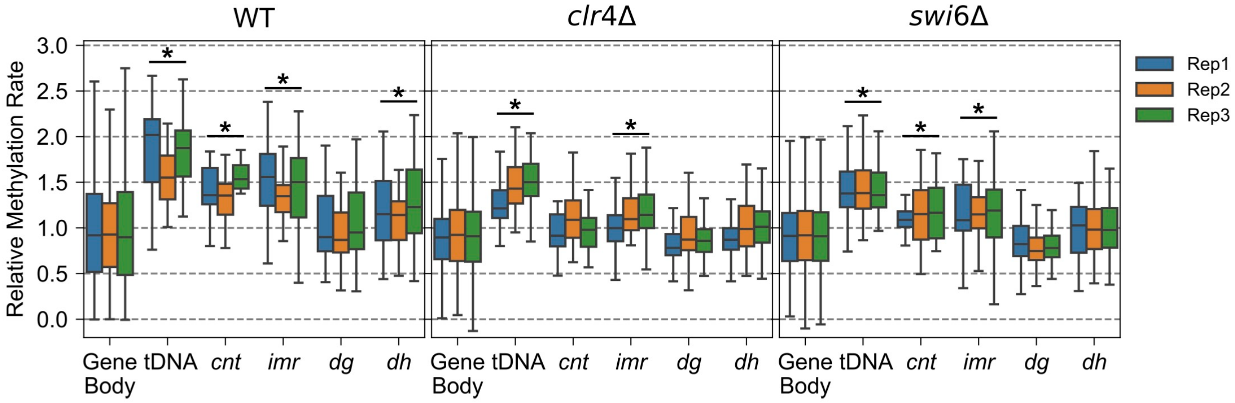


**Supplementary Fig. S4.** Relative methylation rate analysis for WT, *clr4Δ* and *swi6Δ* cells: Comparison of biological replicate experiments. Box plots showing the distribution of methylation rate constants for GATC sites in different genomic regions. Each box contains 25 to 75% of the data, the central line is the median, and the whiskers represent 1.5 times the interquartile range to the farthest data points. Asterisk indicates that the feature is significantly different from gene bodies (Mann-Whitney U test). See Fig. 3D for box plots showing the average methylation rates for the three biological replicate experiments shown here.

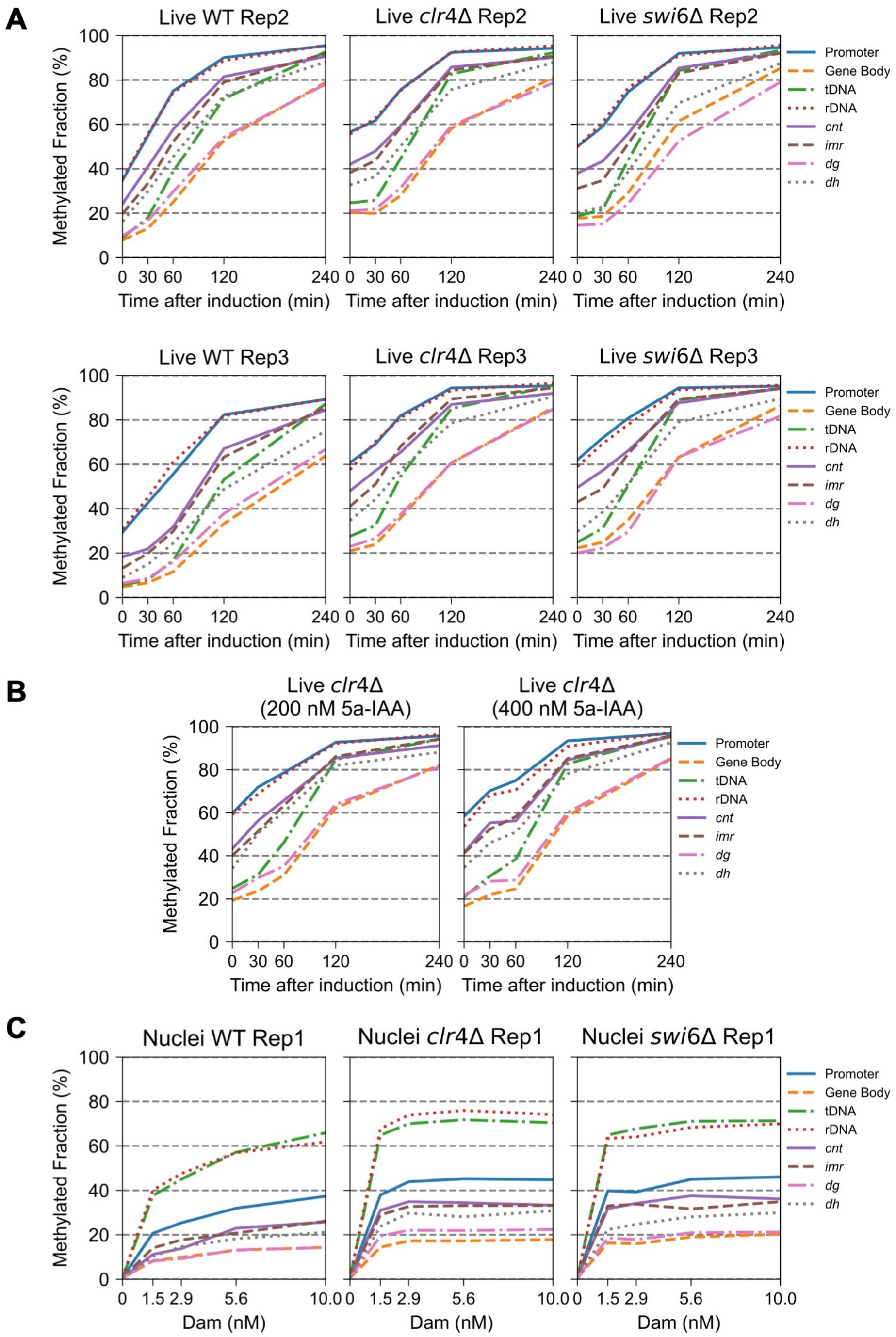


**Supplementary Fig. S5.** Methylation profiles in vivo and in nuclei for various genomic regions: Biological replicate experiments. (**A**) Dam methylation time courses showing the median rate for various genomic regions in vivo using 100 nM 5a-IAA prior to induction (see Fig. 3B for Replicate 1). (**B**) Dam methylation time courses showing the median rate for various genomic regions in *clr4Δ* cells using 200 nM or 400 nM 5a-IAA prior to induction. (**C**) Median methylation of various genomic regions as a function of Dam concentration in wild type, *clr4Δ* and *swi6Δ* nuclei (see Fig. 5C for Replicate 2).

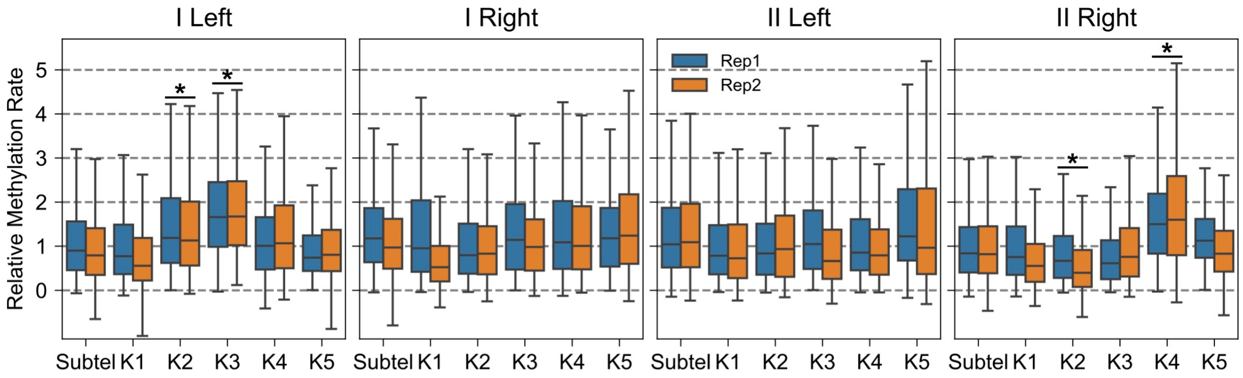


**Supplementary Fig. S6.** Condensed knob chromatin is accessible in vivo. M.SssI methylation rates relative to the genomic median rate for the sub-telomeric regions and the knob regions of chromosomes I and II. Each knob region was divided into five 10-kb windows beginning at the end of the sub-telomeric region as defined by the absence of H3K9me. Each box contains 25 to 75% of the data, the central line is the median, and the whiskers represent 1.5 times the interquartile range to the farthest data points. Asterisk indicates that the feature is significantly different from gene bodies (Mann-Whitney U test).

**Supplementary Table S1.** ***S. pombe*** **strains used in this study.**

| Strains | Genotype | Source |
|---|---|---|
| **972h-** | h- | ATCC 24843 |
| **SP1L** | h- *urg1::ACE1* Dam-degron-3HA kanMX3 | This study |
| **SP3L** | h- *urg1::ACE1* Dam-degron-3HA kanMX3<br>*arg3*::hph1 OsTIR1 (F74A) | This study |
| **SPT999a** | *Mat1-Msmt0 leu1-32 ade6-210 his2⁻ ura4DS/E otr1R::ura4⁺* | Shiv Grewal Lab |
| **SP12L** | *Mat1-Msmt0 leu1-32 ade6-210 his2⁻ ura4DS/E otr1R::ura4⁺*<br>*arg3*::hph1 OsTIR1 (F74A) | This study |
| **SP15L** | *Mat1-Msmt0 lue1-32 ade6-210 his2⁻ ura4DS/E otr1R::ura4⁺*<br>*urg1::ACE1* M.SssI-degron-3HA kanMX3<br>*arg3*::hph1 OsTIR1 (F74A) | This study |
| **SP23L** | h- *urg1::ACE1* Dam-degron-3HA kanMX3<br>*arg3*::hph1 OsTIR1 (F74A) *clr4*::NatNT2 | This study |
| **SP24L** | h- *urg1::ACE1* Dam-degron-3HA kanMX3<br>*arg3*::hph1 OsTIR1 (F74A) *swi6*::NatNT2 | This study |
| **SP27L** | h- *urg1::ACE1* Dam-degron-3HA kanMX3<br>*gcn5-3HA*::NatNT2 arg3::hph1 OsTIR1 (F74A) | This study |
| **SP32L** | h- *clr4*::NatNT2 | This study |
| **SP33L** | h- *swi6*::NatNT2 | This study |
